# Ataxin-2 preserves oocyte genome integrity by promoting timely premeiotic DNA replication

**DOI:** 10.1101/2025.06.13.659559

**Authors:** Vernon L. Monteiro, Crystal Yu, Camilla Roselli, Baskar Bakthavachalu, Mani Ramaswami, Thomas R. Hurd

## Abstract

The faithful reassortment and transmission of chromosomes across generations is fundamental to species survival. While much is known about chromosome pairing and recombination, the upstream regulators controlling entry into the meiotic program remain largely elusive. In many species, including Drosophila and mammals, the decision to enter meiosis occurs prior to premeiotic DNA replication and is governed by post-transcriptional regulation, by yet to be discovered factors. Here, we identify the RNA-binding protein Ataxin-2 as a crucial and previously unrecognized regulator of meiotic entry. We show that Ataxin-2 acts post-transcriptionally to promote the entry into meiosis by downregulating the conserved cell cycle inhibitor Dacapo, the *Drosophila* ortholog of p21/p27. In the absence of Ataxin-2, germ cells mis-regulate Dacapo leading to delayed premeiotic DNA replication and sterility. Strikingly, when DNA replication is delayed and extends into the next stage of meiosis, synaptonemal complex formation, oocytes incur severe genomic DNA damage, likely caused by collisions between replication forks and the synaptonemal complex. Our findings establish Ataxin-2 as a pivotal factor in regulating premeiotic DNA replication and safeguarding oocyte genome stability, shedding new light on the intricate regulatory mechanisms that ensure successful meiosis and fertility.

## INTRODUCTION

Meiosis is essential for gamete production to ensure faithful transmission of genetic information across generations. While it shares some mechanisms and molecular players with mitosis, meiosis employs distinct regulatory programs that enable germ cells to accurately segregate half of each chromosome pair into four gametes^1–4^. To achieve this, germ cells must transition from a mitotic to a meiotic program, beginning with a single round of premeiotic DNA replication followed by two successive nuclear divisions. This premeiotic DNA replication is critical for generating the necessary chromosome complement, ensuring that each gamete inherits a complete set of genetic marterial^4–6^.

Across all studied organisms, premeiotic DNA replication takes significantly longer to complete than mitotic DNA replication and involves distinct regulatory mechanisms^7–12^. While the reason for this remains poorly understand, studies in *Saccharomyces cerevisiae* suggest that the extended premeiotic S-phase aids in the preparation of meiotic recombination^10,13^. Notably, disrupting replication fork stability during premeiotic DNA replication in *Schizosaccharomyces pombe* results in altered meiotic recombination and chromosome missegregation^14^. Yet, despite its fundamental role, the regulation of premeiotic DNA replication in metazoans remains poorly understood.

*Drosophila* and mammals share key aspects of germ cell development. In both *Drosophila* and mammals, germ cells undergo multiple rounds of mitotic division with incomplete cytokinesis before transitioning into meiosis^15,16^. Within these interconnected cells, known as cysts or nests in mammals, only a subset of the germline cells will ultimately develop into the oocyte^17–19^. However, all cyst cells initially transition from mitosis to meiosis, undergoing premeiotic DNA replication roughly concurrently^9^. As meiosis progresses, subsequent meiotic steps such as synaptonemal complex (SC) formation, generation of DNA double-stranded breaks (DSBs) and meiotic recombination become increasingly restricted to only the cells that will become the future oocyte^9,16,17,20^. The remaining cells transfer cytoplasmic content into cells destined to be the oocyte before ultimately dying^17,19,21^. Despite its critical role in fertility, the molecular mechanisms that trigger this initial transition from mitosis to meiosis remain largely unknown.

Recent transcriptome profiling studies have begun to shed light on this transition, revealing that many meiotic genes are transcribed before meiosis begins but remain untranslated^22,23^. This suggests that RNA-binding proteins (RBPs) are required to regulate these mRNAs post-transcriptionally, promoting their translation and driving meiotic entry and progression. While RBPs have been implicated in suppressing mitosis^24–27^, the key RBPs responsible for promoting mitotic to meiotic transition beginning with premeiotic DNA replication remain unidentified. Identifying these factors is crucial for understanding the molecular switch that propels germ cells into meiosis.

We recently discovered that the RBP Ataxin-2 (Atx2) functions precisely at the stage when germ cells transition into meiosis^28^, positioning it as a potential key regulator of meiotic entry. Consistent with this, in both *Drosophila* and *Caenorhabditis elegans*, Atx2 is essential for oogenesis, underscoring its conserved role in germ cell development^29–31^. Furthermore, Atx2 typically enhances the stability and translation of its target mRNAs by recruiting polyadenylation complexes and interacting with poly(A)-binding proteins (PABPs)^32–37^, making it ideally suited to function as a positive regulator of meiotic entry.

Here, we show that in *Drosophila*, Atx2 regulates the timing of premeiotic DNA replication. We find that Atx2 promotes the downregulation of the conserved cell cycle inhibitor p21/p27, known as Dacapo (Dap) in *Drosophila*. In *Atx2*-depleted female flies, Dap fails to be downregulated, leading to delayed entry into premeiotic S-phase and sterility. Although germ cells eventually undergo premeiotic DNA replication, they do so in a delayed and unsynchronized manner, resulting in irreparable genomic damage specifically in the oocyte. Our genetic data suggest that this damage arises from replication stress associated with delayed DNA replication overlapping with SC formation. Together, our findings establish Atx2 as a critical regulator of meiotic entry and an essential guardian of oocyte genome stability in Drosophila. Remarkably, recent discoveries in mammals show that timely entry into premeiotic DNA replication is likewise crucial for preserving oocyte genome integrity^38^, underscoring the fundamental importance of this regulatory mechanism across species.

## RESULTS

### Loss of Atx2 activates the meiotic checkpoint

Through a previous large-scale RNAi screen, we uncovered an unexpected role for Atx2 at the onset of meiosis^28^, raising the intriguing possibility that Atx2 serves as a broader regulator of meiotic entry and progression. To test this, we depleted *Atx2* in the germline using a pan-germline driver (MTD-GAL4)^39^ and observed a near-complete abolishment of embryo hatching with two independent RNAi strains (Fig. 1A) and a minor reduction in egg laying (Fig. S1A). To investigate the hatching defect, we examined embryos from *Atx2*-depleted females and observed eggshell abnormalities. Unlike wild-type embryos, which form two dorsal appendages, the breathing tubes of the embryo, *Atx2* knockdown embryos often had a single, thin or fused dorsal appendage, with some embryos lacking them altogether (Fig. 1B, C). This classic ‘spindle’ phenotype is typically caused by reduced levels of the protein Gurken, a TGFα-like ligand essential for dorsal-ventral patterning in the egg and antero-posterior axis formation in the embryo^40^ (Fig. 1D). A frequent cause of Gurken reduction is activation of the meiotic checkpoint^41^ (Fig. 1D). Thus, these findings show that Atx2 is critical for fertility possibility by regulating meiotic progression.

**Figure 1.**
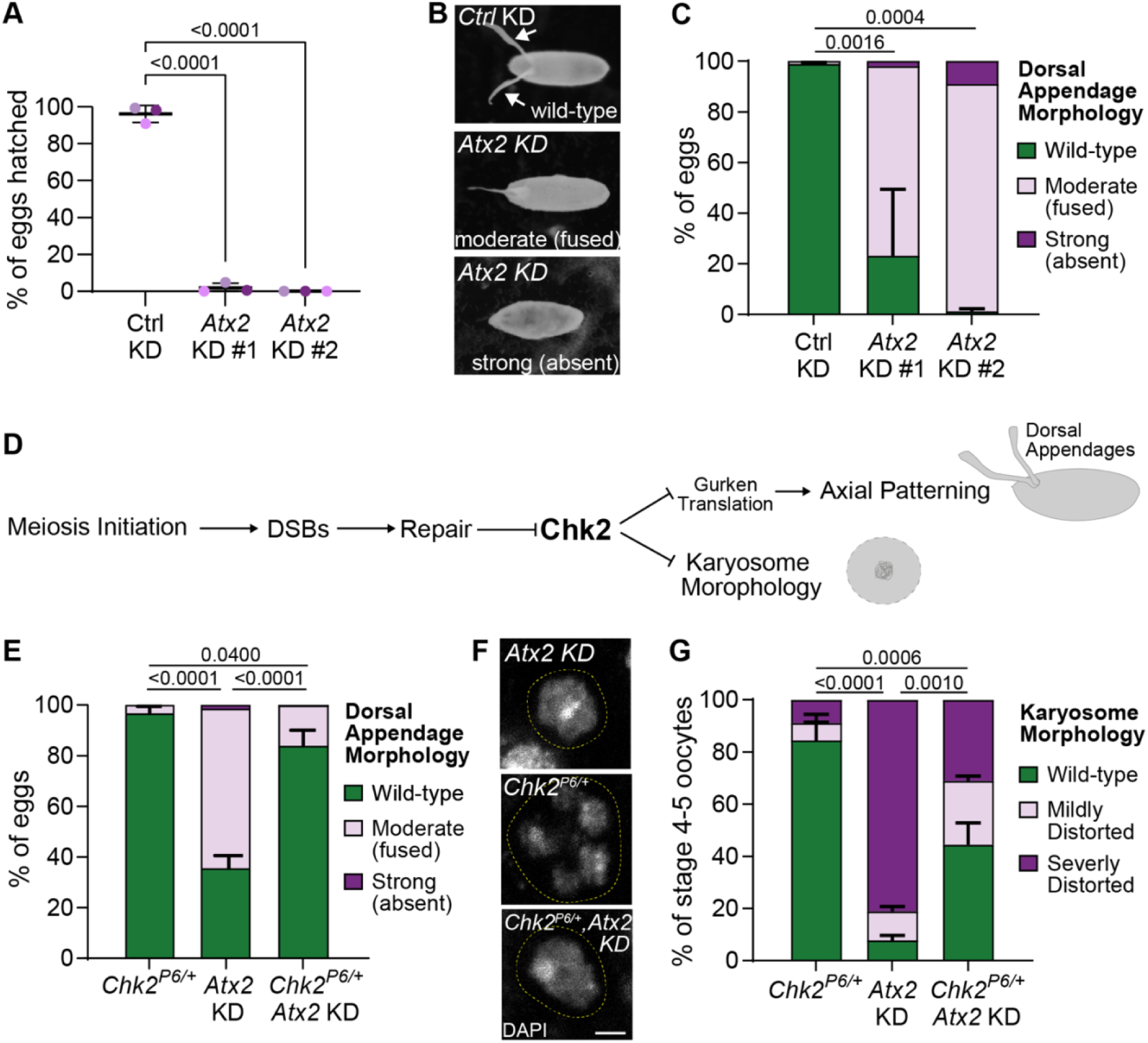
Atx2 depletion activates the meiotic checkpoint. **(A)** Percent of eggs from females expressing Ctrl (*mCherry*) or *Atx2* shRNA that hatched. Data are means ± s.d. of three independent replicates (>100 eggs per replicate). *p*-values, one-way ANOVA followed by Dunnett’s multiple comparison test. **(B)** Images of eggshells from females expressing Ctrl (*mCherry*) shRNA or *Atx2* shRNA. **(C)** Quantification of (**B**). Percent of eggs from females of the indicated genotypes with wild-type, fused or strong dorsal appendage morphology phenotypes. Data are means ± s.d. of three independent replicates (100 – 831 eggs per replicate). *p*-values, one-way ANOVA followed by Dunnett’s multiple comparison test on the wild-type eggshell category. **(D)** Cartoon of the meiotic checkpoint showing that a failure to repair DNA damage activates Chk2 impairing both axial patterning (by inhibiting Gurken translation) and karyosome morphology. **(E)** Percent of eggs from females of the indicated genotypes with wild-type, fused or strong dorsal appendage morphology phenotypes. Data are means ± s.d. of three independent replicates (231 – 1080 eggs per replicate). *p*-value, ANOVA with Tukey’s multiple comparison test on wild-type eggshell category. **(F)** Karyosome (dashed yellow line) morphology of stage 4-5 oocytes of indicated genotypes. Scale bar, 2 µm. **(G)** Quantification of **(F)**. Mildly distorted (loss of sphericity, condensed); severely distorted (uncondensed). Data are means ± s.d. of three independent replicates (30 oocytes per replicate). *p*-values, ANOVA with Tukey’s multiple comparison test on spherical karyosomes.

*Checkpoint kinase 2* (Chk2) is the master regulator of the meiotic checkpoint. When activated during oogenesis it prevents Gurken translation causing the spindle phenotype and disrupting karyosome morphology^41,42^ (Fig. 1D). Loss of just one copy of *Chk2* suppresses checkpoint activation and can partially restore both phenotypes^42^. Thus, to determine whether the observed eggshell defects in *Atx2* depletion embryos are due to activation of the meiotic checkpoint, we knocked down *Atx2* in a background *Chk2* heterozygous null mutant. As before, we found that silencing *Atx2* in the germline using a *nos*-GAL4 driver, which is expressed most highly at the onset of meiosis, caused a strong spindle phenotype (Fig. 1E) and a reduction in oocyte Gurken protein (Fig. S1B, C). This was caused by activation of the meiotic checkpoint, as loss of one functional copy of *Chk2* largely restored dorsal appendage morphology (Fig. 1E). To confirm that *Atx2* depletion triggers checkpoint activation, we also examined oocyte karyosome morphology. As expected, we observed abnormal oocyte karyosome morphology in *Atx2* knockdowns (Fig. 1F, G), which was partially rescued by removing one copy of *Chk2* (Fig. 1F, G). Together, these findings indicate that depletion of *Atx2* during meiosis triggers activation of the meiotic checkpoint.

### *Atx2* maintains oocyte genome stability during meiosis

To investigate Atx2’s role in meiosis, we examined the germarium—the anterior region of the ovary where meiosis is initiated (Fig. 2A). In region 1, oogenesis begins when a germline stem cell divides asymmetrically to produce a cell that undergoes four mitotic divisions with incomplete cytokinesis to form a 16-cell cyst. Meiosis immediately follows in region 2, with all 16 cyst cells first synchronously undergoing premeiotic DNA replication, after which SC formation, double-strand break (DSB) induction, and recombination become progressively restricted to a single cyst cell^3,9^. By region 3, only one cell—the oocyte—remains in meiosis, having completed recombination and repaired its DSBs, while the other 15 have become nurse cells that support oocyte development^18^. If DSBs are not fully repaired by region 3, the meiotic checkpoint is triggered, resulting in eggshell defects, abnormal karyosomes, and ultimately sterility^42–44^.

**Figure 2.**
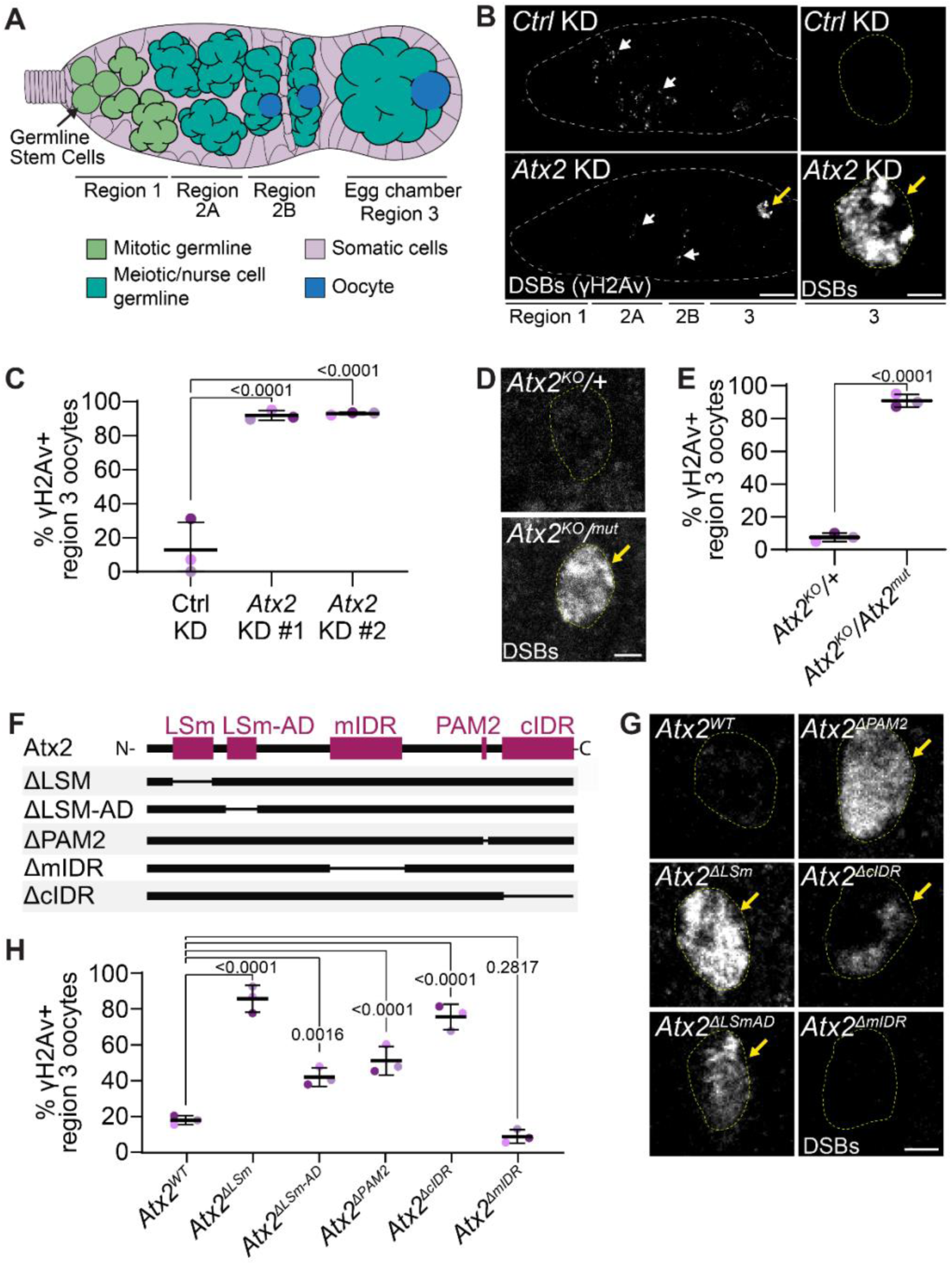
*Atx2* maintains oocyte genome stability during meiosis. (A) Schematic of female Drosophila germarium subdivided into regions 1, 2A and 2B, and 3 based on developmental stages. (B) Confocal micrographs of germaria (left) and region 3 oocytes (right) expressing Ctrl (*mCherry*) or *Atx2* shRNA in the germline stained with anti-γH2Av. Yellow arrows indicate persistent γH2Av staining. White arrows indicate meiotic γH2Av staining. Scale bars, 10 µm (germarium) and 2 µm (oocyte). (C) Frequency of γH2Av positive region 3 oocytes expressing Ctrl (*mCherry*) or *Atx2* shRNAs. Data are means ± s.d. of three independent replicates (30 – 100 oocyte per replicate). *p*-values, ANOVA, Dunnett’s multiple comparison test. (D) Confocal micrographs of region 3 oocytes from heterozygous control and germline ablated *Atx2* ovaries stained with γH2Av. Scale bars, 2 µm (oocyte). (E) Frequency of γH2Av positive region 3 oocytes expressing Ctrl (*mCherry*) or *Atx2* shRNAs. Data are means ± s.d. of three independent replicates (13 – 40 oocytes per replicate). *p*-values, two-tailed unpaired t-test. (F) Schematic of Atx2 protein domains and domain deletion mutants. (G) Confocal micrographs of region 3 oocytes expressing full-length *Atx2*, *Atx2^ΔLSm^*, *Atx2^ΔLSm-AD^*, *Atx2^ΔPAM2^*, *Atx2^ΔmIDR^* and *Atx2^ΔcIDR^* under native regulatory elements in *Atx2* germline ablated ovaries stained with anti-γH2Av. Scale bars, 2 µm. (H) Frequency of γH2Av positive region 3 oocytes of the indicated genotypes. Data are means ± s.d. (24 – 40 oocytes per replicate). *p*-values, ANOVA, Dunnett’s multiple comparison test.

To test whether Atx2 is required to maintain oocyte DNA integrity or repair DSBs during meiosis, we assessed DSB persistence in region 3 oocytes using an antibody against the phosphorylated histone variant H2Av (γH2Av), the Drosophila homolog of both H2Az and H2Ax^45–47^. In control and *Atx2* knockdown ovaries, γH2Av forms distinct foci in the 16-cell cysts of region 2, marking the programmed DSBs that occur during normal meiotic recombination^47,48^ (Fig. 2B, white arrows). In control ovaries, these DSBs are repaired as meiosis proceeds and by region 3, γH2Av staining is not present in oocytes (Fig. 2B, C, S2A). In contrast, *Atx2* knockdowns γH2Av staining persisted into region 3 at high frequencies (Fig. 2B, C, S2A). Rather than discrete foci, the γH2Av signal often blanketed the entire oocyte chromatin (Fig. 2B), consistent with extensive unrepaired DNA damage. Strikingly, γH2Av signal was confined to the oocyte and was not detected in the mitotically dividing germ cells of region 1 or strongly in the nurse cells of region 3 above that of wildtype levels, indicating a specific requirement for Atx2 in protecting oocyte genomic integrity (Fig. 2B, S2A).

To validate our findings and rule out off-target effects, we generated germline-specific *Atx2* knockouts using an FRT-flanked *Atx2* allele and we also conducted genetic rescue experiments. Consistent with our knockdown results, we detected strong γH2Av signal in region 3 oocytes of *Atx2* germline knockouts, but not in controls (Fig. 2D, E, S2B). Similarly, expression of an shRNA resistant wildtype *Atx2* restored genome integrity in the *Atx2* knockdown background (Fig. S2C, D). Notably, of the two *Atx2* isoforms expressed in the ovary^49–51^, only *Atx2-RB*, which encodes an additional 61 amino acid N-terminal extension of unknown function, fully rescued the DNA damage phenotype indicating a specific requirement for the RB isoform (Fig. S2C, D). These results establish a critical and oocyte-specific role for Atx2 in maintaining genome stability during meiosis.

To understand how Atx2 promotes oocyte genome stability, we examined its functional domains. Atx2 contains several conserved features (Fig. 2F): the Like-Sm (LSm) and LSm-associated domain (LSm-AD), both implicated in mRNA binding and processing^36^; the PABP interaction motif (PAM2), essential for binding Poly(A)-binding protein (PABP)^36^; and two intrinsically disordered regions (IDRs)—the middle IDR (mIDR) and C-terminal IDR (cIDR)—which promote ribonucleoprotein (RNP) granule formation^52^. The cIDR has recently also been implicated in binding mRNAs^53^, leaving the mIDR as the only region uniquely associated with granule formation. To determine which domains are essential for oocyte genome maintenance, we expressed *Atx2* deletion mutants in a germline-specific *Atx2* knockout background^29,52^. Expression of full-length *Atx2* effectively restored oocyte genome integrity in the *Atx2* knockout background (Fig. 2G, H, S2E), while deletion versions lacking the LSm, LSm-AD, PAM2, or cIDR regions failed to do so (Fig. 2G, H, S2E). Interestingly, expression of *Atx2* lacking its mIDR restored genome stability to wild-type levels (Fig. 2G, H, S2E). This indicates that domains involved in mRNA regulation, but not those solely associated with RNP granule formation are required for maintaining oocyte genome stability. This suggests that Atx2 safeguards genome stability through its role in post-transcriptional gene regulation.

### Atx2 maintains genome stability independently of DSB repair and transposon repression

Given that Atx2 preserves oocyte genome stability through post-transcriptional gene regulation, we considered whether it does so by promoting meiotic DSB repair, potentially through the upregulation of DNA repair genes. To investigate this, we blocked programmed meiotic DSB formation using loss-of-function mutants in *mei-W68* (the Drosophila homolog of *SPO11*) and *mei-P22* (a TopoVIB-like homolog)^54–56^. If genome instability in *Atx2*-depleted oocytes is due to a lack of DSB repair, preventing programmed DSB formation altogether should rescue the damage phenotype. As expected, γH2Av foci, indicative of DSBs, were absent in region 3 of control and *mei-P22* (Fig. S3A, B)^47^. However, strikingly, strong γH2Av signal persisted in *Atx2*-depleted oocytes even when programmed meiotic DSBs were abrogated by mutating either *mei-P22* or *mei-W68* (Fig. S3A, B). These results reveal that Atx2 does not promote meiotic DSB repair. Instead, it appears to safeguard the oocyte genome through an alternative, DSB-repair independent mechanism.

De-repression of transposons contributes to genomic instability in the Drosophila germline^57–60^ and is known to activate the meiotic checkpoint^59^. As *Atx2* was previously shown to regulate micro-RNAs *in vivo*^61^, we wondered if Atx2 might instead maintain oocyte genome integrity by repressing transposons through another small RNA pathway, the piRNA pathway^60^. Thus, we measured the levels of two abundant retrotransposons *blood* and *burdock* by RT-qPCR^58,62,63^. Knockdown of *aubergine* (*aub*), an essential PIWI-family protein which represses transposons^64^, resulted in dramatic increase of *blood* and *burdock* transcripts (Fig. S3C, D). However, *Atx2* knockdown did not result in their increase compared to the control knockdown (Fig. S3C, D), indicating that the genomic instability observed in *Atx2* depletion is not a result of increased transposon activity. Together, our data indicate that Atx2 maintains oocyte genome stability through a mechanism independent of both DSB repair and transposon repression.

### Atx2 regulates timely premeiotic DNA replication

To uncover how Atx2 maintains oocyte genome integrity during meiosis, we next asked whether it ensures proper execution of DNA replication at this decisive developmental stage. Disruptions to DNA replication can lead to replication stress, a catastrophic event that threatens genome stability and activates DNA damage responses^65^. Thus, we measured how *Atx2* knockdown influences premeiotic DNA replication using 5-ethynyl-2’-deoxyuridine (EdU) incorporation and confocal microscopy^66^. In wild-type germaria, we captured EdU incorporation in all 16 cells of newly formed cysts in region 2A, reflecting the synchronous onset of premeiotic DNA replication^9^, in ∼40% of the germarium analyzed (Fig. 3A yellow stars and 3B). In stark contrast, in *Atx2* knockdown germaria, we observed a significant reduction in number of 16-cell cysts with uniform EdU incorporation in region 2A (Fig. 3A yellow stars and 3B), alongside an increase in EdU labeling in region 2B (Fig. 3A white arrows). Notably, this delayed replication in region 2B of *Atx2* knockdown cysts often appeared asynchronous, with only a subset of cells within the cyst incorporating EdU (Fig. 3A white arrows). Germline-specific deletion of *Atx2* produced a similar defect: synchronous premeiotic DNA replication was delayed, and EdU signal was often detected in only a few cells per cyst in region 2B, with many cysts showing incomplete incorporation, often with fewer than half the cells labeled (Fig. 3C–E)—a phenotype never observed in controls. These results uncover a previously unappreciated function for Atx2 in coordinating the onset of DNA replication at meiosis. Loss of *Atx2* disrupts this process, leading to delayed and asynchronous premeiotic DNA replication—an aberration likely to drive the observed genome instability (Fig. 2) and checkpoint activation (Fig. 1).

**Figure 3.**
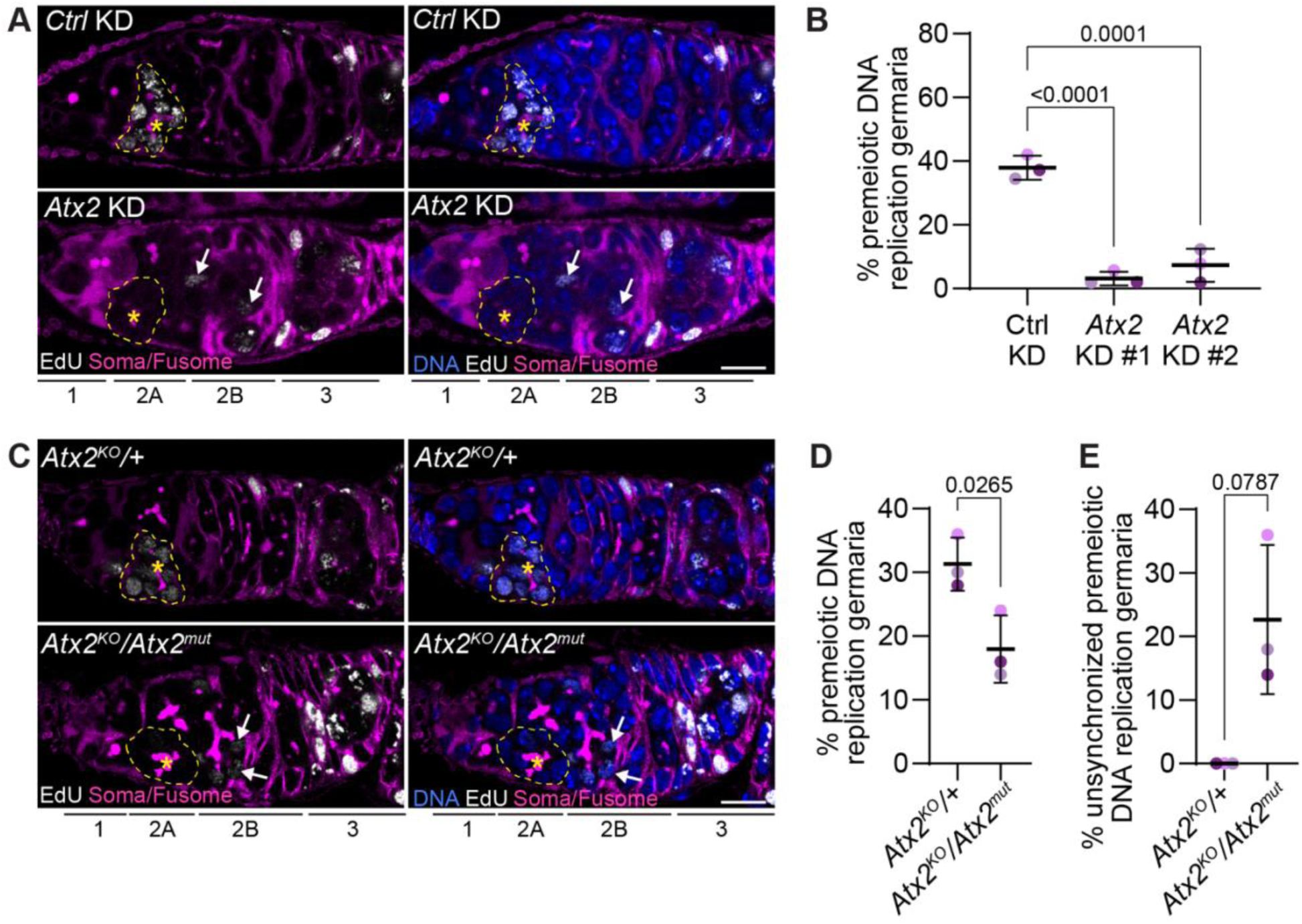
Premeiotic DNA replication is delayed and unsynchronized in the absence of *Atx2*. **(A)** Confocal micrographs of germaria expressing Ctrl (*mCherry*) or *Atx2* shRNA in the germline stained with EdU (white), anti-1B1 (soma/fusome, magenta) and DAPI (DNA, blue). Scale bar, 10 µm. Yellow dashed line, early region 2A cyst. **(B)** Quantification of **(C)**. Percent germaria with 16-cell cysts undergoing premeiotic DNA replication from indicated genotypes. Data are means ± s.d. (43 – 57 germaria per replicate). *p*-value, ANOVA, Dunnett’s multiple comparison test. **(C)** Confocal micrographs of heterozygous control and Atx2 germline ablated germaria stained with EdU (white), anti-1B1 (soma/fusome, magenta) and DAPI (DNA, blue). Scale bar, 10 µm. Yellow dashed line, early region 2A cyst. Scale bar, 10 µm. **(D, E)** Quantification of **(C)**. Percent of germaria with 16-cell cysts undergoing premeiotic DNA replication **(D)** and those with unsynchronized premeiotic DNA replication (**E**). Data are means ± s.d. (50 germaria per replicate). *p*-values, two-tailed unpaired t-test **(D)** and two-tailed Welch’s t-test **(E)**.

### Atx2 promotes timely premeiotic DNA replication by downregulating the cyclin-dependent kinase inhibitor Dacapo

We next sought to elucidate the mechanism by which Atx2 controls premeiotic DNA replication. We first focused on the cyclin-dependent kinase inhibitor Dacapo (Dap)—a pivotal cell cycle regulator previously implicated in regulating premeiotic DNA replication in the Drosophila ovary^67^. Dap inhibits cyclin-CDK activity to allow assembly of the pre-replication complex but must be downregulated to initiate DNA replication^67–70^. Notably, in the ovary Dap is specifically required for premeiotic DNA replication in the 16-cell cysts, but not for the preceding mitotic cycles^67^. We therefore hypothesized that Atx2 promotes premeiotic replication by driving the timely downregulation of Dap.

Using an endogenously tagged Dap-GFP reporter^71^, we examined Dap protein levels in region 2A. In wild-type germaria, we detected Dap-GFP signal ∼20% of early region 2A 16-cell cysts, representing cysts that had recently completed mitosis but had not yet initiated premeiotic DNA replication (Fig. 4A yellow star; 4B early 2A). By late region 2A, Dap-GFP was largely absent, indicating the onset of DNA replication (Fig. 4A white star; 4B late 2A), but reappeared in late region 2B and region 3, where Dap is known to regulate nurse cell endocycling and maintain oocyte fate^68,72^. Strikingly, *Atx2* knockdown germaria showed sustained and elevated Dap protein across both early and late region 2A (Fig. 4A, B), indicating a failure to properly downregulate Dap. We further examined Cyclin E (CycE), which is bound and stabilized by Dap, thereby inhibiting its function^69,70^, using a CycE-GFP reporter. Consistent with elevated Dap levels, CycE also persisted abnormally in *Atx2*-depleted germaria (Fig. S4A, B), reinforcing the conclusion that Atx2 facilitates the transition into DNA replication by reducing Dap protein levels.

**Figure 4.**
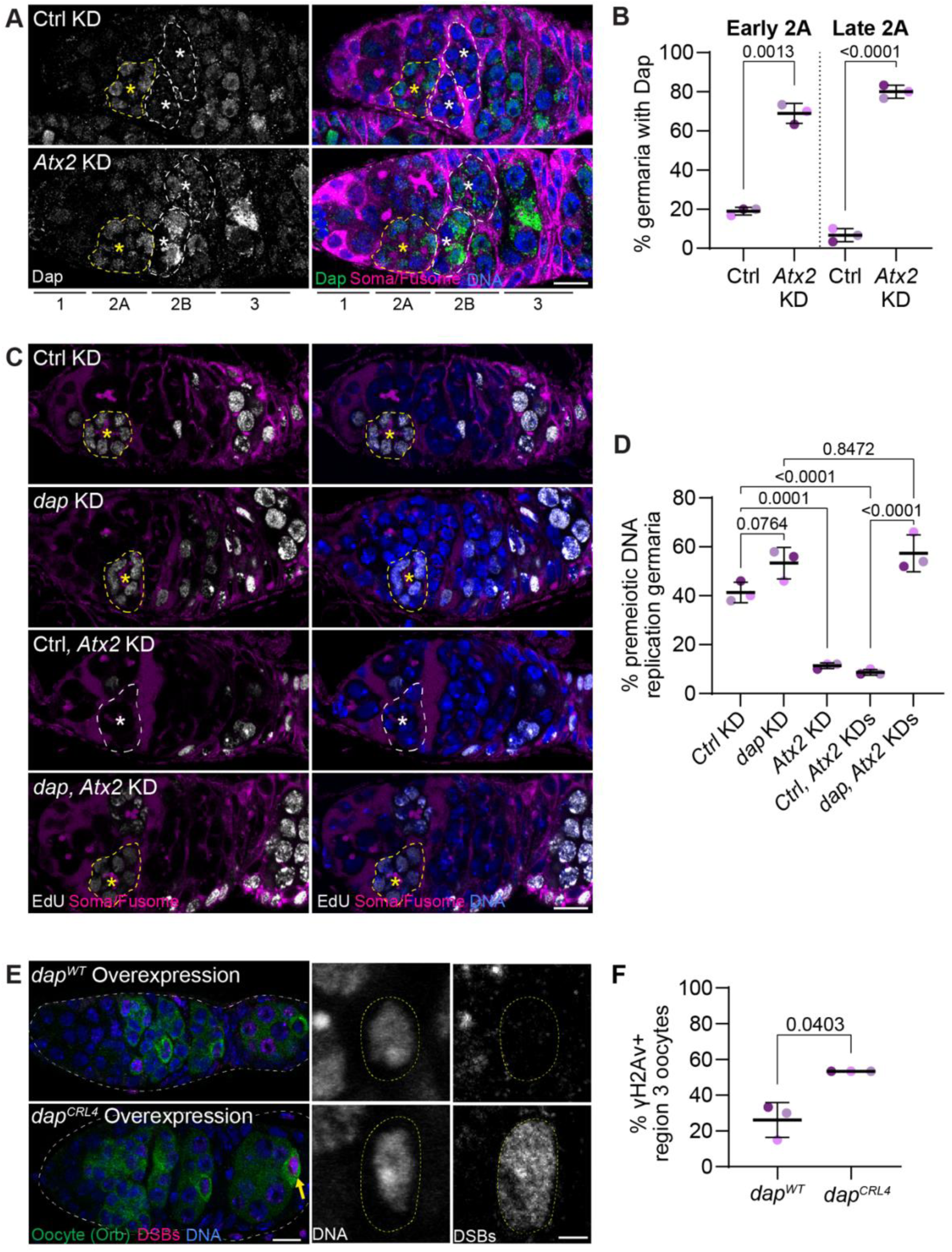
Atx2 regulates Dap to preserve oocyte genomic integrity. **(A)** Confocal micrographs of Dap-GFP germaria expressing Ctrl (*LexA*) or *Atx2* shRNAs in the germline stained with anti-GFP (Dap, green), anti-1B1 (soma/fusome, magenta) and DAPI (DNA, blue). Scale bar, 10 µm. Yellow dashed line, early region 2A. While dashed line, region 2B. **(B)** Quantification of **(A)**. Percent of germaria with above background levels of Dap-GFP in the early or late region 2A cysts of the indicated genotypes. Data are means ± s.d. (25 – 30 germaria per replicate). *p*-value, two-tailed Welch’s t test. **(B)** Premeiotic DNA replication in germaria expressing **(D)** Ctrl (*LexA*), **(E)** *dap*, **(F)** Ctrl and *Atx2*, and **(G)** *dap* and *Atx2* shRNAs under the control of *nos-GAL4.* Scale bar, 10 µm. Yellow dashed line, region 2A cyst. **(C)** Quantification of **(C)**. Percent of germaria with 16-cell cysts undergoing premeiotic DNA replication of the indicated genotypes. Data are means ± s.d. (50 germaria per replicate). *p*-value, ANOVA, Tukey multiple comparison test. **(D)** Confocal micrographs of germaria (left) and region 3 oocytes (right) expressing full-length *dap* or *dap^CRL4^* in the germline stained with anti-γH2Av (DSB, magenta), anti-Orb (oocyte, green) and DAPI (DNA, blue). Scale bar, 10 µm (germaria) and 2 µm (oocyte). White dashed line, germline. Yellow dashed line, oocyte nucleus. **(E)** Quantification of **(E)**. Percentage of γH2Av positive region 3 oocytes from indicated genotypes. Data are means ± s.d. (20 – 30 oocytes per replicate). *p*-values, two-tailed Welch’s t test.

To determine whether elevated Dap levels are functionally responsible for the replication delay observed in *Atx2* mutants, we conducted genetic epistasis experiments. While *dap* knockdown alone caused only a modest increase in the number of 16-cell cysts undergoing premeiotic replication (Fig. 4C, D), strikingly, co-depletion of both *Atx2* and *dap* completely restored replication frequency to the level seen with *dap* knockdown alone. This finding demonstrates that elevated Dap is the primary barrier to premeiotic DNA replication in *Atx2* mutants and that reducing Dap is sufficient to bypass the requirement for Atx2. These results uncover a previously unrecognized role for Atx2 in promoting premeiotic DNA replication by repressing Dap, revealing a critical post-transcriptional mechanism that ensures timely cell cycle progression at the onset of meiosis.

### Delaying premeiotic DNA replication triggers oocyte genome instability

It remained unclear whether the genomic instability seen in *Atx2* depletion stems primary from a failure to properly downregulate Dap and initiate premeiotic DNA replication at the correct time. To resolve this, we asked whether simply blocking Dap downregulation induces DNA damage in the oocyte. To do this, we overexpressed wild-type Dap as a control and a degradation-resistant form of Dap (Dap^CRL4^), which lacks a known CRL4^l(2)dtl^ degron motif^73^. Notably, region 3 oocytes expressing *dap^CRL4^* exhibited a significantly higher frequency of γH2Av signal compared to those overexpressing wild-type *dap* (Fig. 4E, F), indicating that persistent Dap protein is sufficient to compromise oocyte genome integrity. Together, these findings demonstrate that Atx2 safeguards the oocyte genome by promoting the timely downregulation of Dap and ensuring proper initiation of premeiotic DNA replication.

Finally, to determine whether the oocyte DNA damage reflects a broader consequence of delayed premeiotic DNA replication or is specific to elevated Dap protein, we asked whether disrupting replication through a Dap-independent mechanism would similarly trigger oocyte genomic instability. To delay premeiotic DNA replication, we knocked down *cdc7*, a known regulator of DNA replication initiation^74–76^. Indeed, germline knockdown of *cdc7* led to robust accumulation of γH2Av in oocytes (Fig. S5A, B), consistent with a DNA damage response. Together, these results indicate that delaying premeiotic DNA replication—regardless of the mechanism—can precipitate a catastrophic loss of genome integrity in oocytes.

### Synaptonemal complex aggravates genomic instability in oocyte in absence of Atx2

A striking outcome of delaying premeiotic DNA replication is that DNA damage occurs specifically in the oocyte, even though the replication delay affects all 16 cells within the cyst. One possible explanation lies in replication stress, which can be intensified by conflicts with other cellular processes occurring concurrently with DNA replication^65^. During Drosophila oogenesis, SC assembly begins shortly after premeiotic DNA replication and is maintained only in the pro-oocyte^9,20,77^. Thus, in *Atx2* mutants, we hypothesized that delayed premeiotic DNA replication could result in conflicts with the SC, resulting in genome instability in the oocyte but not the nurse cells. Loss of core SC components such as the transverse filament protein, C(3)G, which connects homologous chromosomes, abolishes formation of the SC and meiotic recombination^77^. To test whether the SC contributes to oocyte-specific DNA damage, we examined γH2Av in *Atx2* knockdown germaria with and without the SC using *c(3)G^1^* null mutants in which the SC is completely absent^77^. Consistent with our hypothesis, γH2Av signal was elevated in region 3 oocytes of *Atx2* knockdown germaria but was rescued to wildtype levels when we removed the SC in this background (Fig. 5A, B). Similarly, depleting *c(3)G* in an *Atx2* knockout background significantly suppressed the DNA damage phenotype (Fig. S6A, B), although to a lesser extent, which may reflect only a partial reduction in the SC due to incomplete gene silencing of *c(3)G*. Together, these results suggest that SC formation exacerbates genome instability in the oocyte.

**Figure 5.**
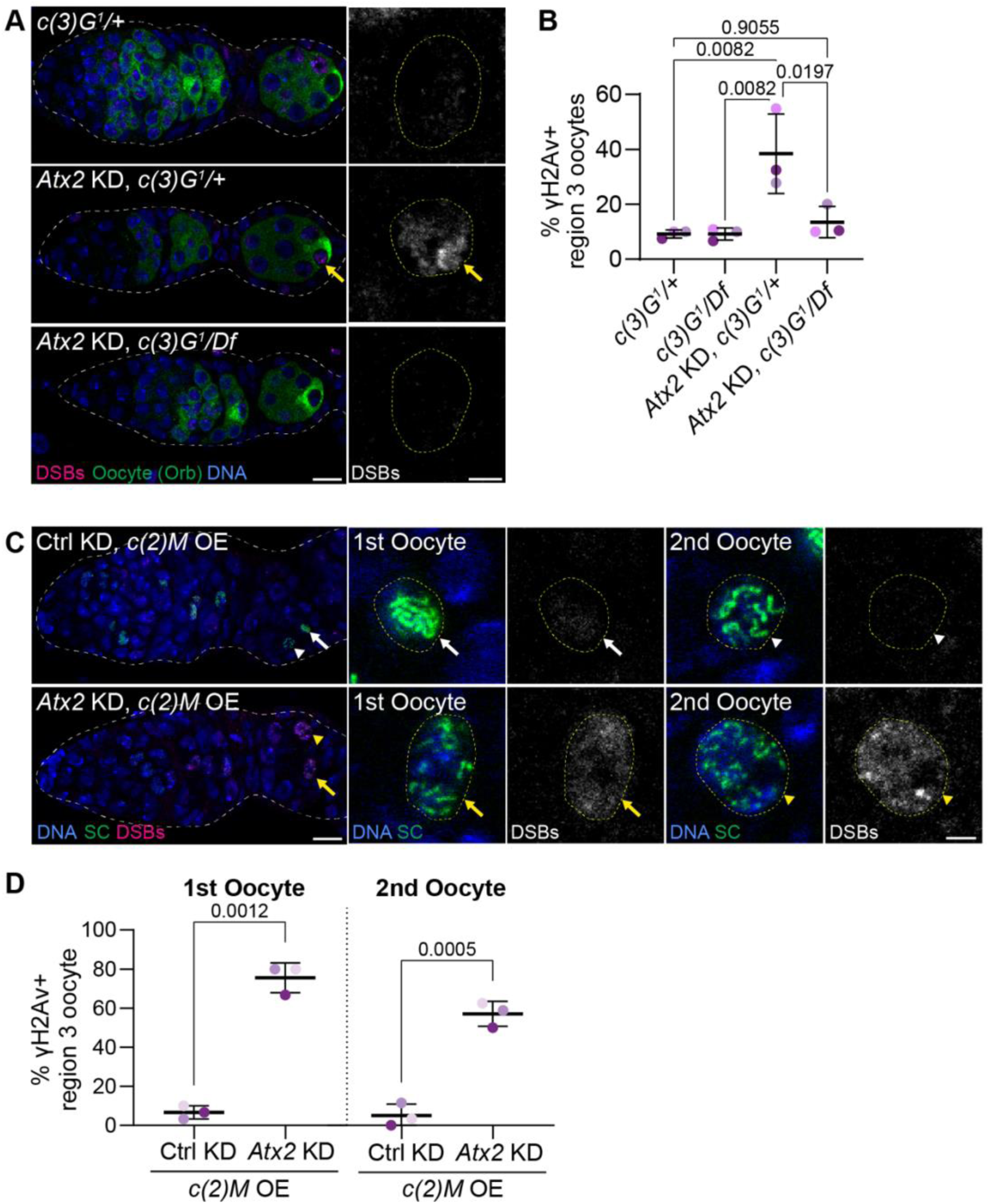
Synaptonemal complex exacerbates oocyte genomic instability. **(A)** Confocal micrographs of germaria (left) and region 3 oocytes (right) of the following genotypes: *c(3)G^1^*/+ heterozygotes; *Atx2* shRNA expressed in the germlines of *c(3)G^1^*/+ heterozygotes; *Atx2* shRNA expressed in the germlines of *c(3)G^1^/Df* transheterozygotes. Ovaries were stained with anti-γH2Av (DSB, magenta), anti-Orb (oocyte, green) and DAPI (DNA, blue). Scale bar, 10 µm (germaria) and 2 µm (oocyte). White dashed line, germline. Yellow dashed line, oocyte nucleus. Yellow arrow, persistent oocyte γH2Av. **(B)** Quantification of **(A)**. Percentage of γH2Av positive region 3 oocytes of the indicated genotypes. Data are means ± s.d. of three independent replicates (18 – 50 oocytes per replicate). *p*-value, ANOVA, Tukey’s multiple comparison test. **(C)** Confocal micrographs of germaria (left) and region 3 oocytes (right) overexpressing *c(2)M* and either Ctrl (*LexA*) or *Atx2* shRNAs in the germline stained with anti-γH2Av (DSB, magenta), anti-Orb (oocyte, green) and DAPI (DNA, blue). Scale bar, 10 µm (germaria) and 2 µm (oocyte). White dashed line, germline. Yellow dashed line, oocyte nucleus. Yellow arrow and arrowheads, persistent oocyte γH2Av in the 1^st^ and 2^nd^ pro-oocyte, respectively. **(D)** Quantification of **(C)**. Percent of γH2Av positive region 3 1^st^ and 2^nd^ pro-oocytes of the indicated genotypes. Data are means ± s.d. of three independent replicates (30 1^st^ and 16–29 2^nd^ pro-oocyte per replicate). *p*-value, Welch’s t-test.

To further test whether the SC itself sensitizes cells to DNA damage in *Atx2* knockdowns, we drove persistent SC formation in the 2nd pro-oocyte into region 3 by overexpressing the lateral element gene *c(2)M* which runs along the chromosome arms^78^. In control knockdowns with *c(2)M* overexpression, the 2nd pro-oocyte retained SC but did not display γH2Av signal (Fig. 5C, D). In contrast, in *Atx2* knockdown germaria overexpressing *c(2)M*, γH2Av was present in both the oocyte and the 2nd pro-oocyte (Fig. 5C, D), indicating that the SC sensitizes these cells to replication-associated DNA damage. Together, these findings demonstrate that the SC contributes to the genome instability triggered by *Atx2* loss and delayed premeiotic replication and highlight the oocyte’s unique vulnerability to DNA damage during early meiosis.

## DISCUSSION

The regulation of meiotic entry remains poorly understood in metazoans. However, a recurring theme across species—and possibly an ancestral mechanism for initiating meiotic protein expression—is the critical role of post-transcriptional regulation. This is especially evident in Drosophila and mice, where many meiotic genes are transcribed before meiotic entry^22,23,79,80^, yet the factors and mechanisms regulating their translation remain largely unknown. Here, we identify the RBP Atx2 as the first such regulator in Drosophila. This finding echoes recent work in mice showing that RBPs YTHDC2 and RBM46, together with MEIOC, coordinate the regulation of specific mRNAs to promote meiotic entry^81–85^.

We show that Atx2 promotes premeiotic DNA replication by repressing the expression of the cell cycle inhibitor Dap. Atx2 is known to regulate RNA stability and translation via its LSm and PAM2 domains^33,34,37,61^, and we find both domains are essential for its function in this context— supporting a role as a translational regulator. While Atx2 may directly bind *dap* mRNA to repress translation, we favor a model in which Atx2 also promotes the expression of proteins that facilitate Dap degradation. Candidates include components of the SCF^Skp2^ and CRL4^Cdt2^ E3 ubiquitin ligase complexes, which are known to target Dap for degradation in other contexts^73,86^. Supporting this, SkpA, a core SCF^Skp2^ component, has recently been implicated in meiotic progress in flies^87^. Our attempts to identify specific Dap targets during meiosis using RNA immunoprecipitation approaches have been challenging. However, determining how Dap is degraded during meiosis is a key next step to understanding its regulation of premeiotic DNA replication in Drosophila.

Strikingly, we found that failure to initiate premeiotic DNA replication at the correct time leads to catastrophic oocyte-specific genomic instability and sterility. Although all 16 germ cells in a cyst exhibit delayed replication, only the oocyte accumulates DNA damage—a phenomenon not previously reported in Drosophila. This specificity likely arises from SC dynamics, which are uniquely maintained in the oocyte^9,20,77^. Under normal conditions, premeiotic DNA replication concludes before full SC assembly^9^. However, in *Atx2* depletions, this temporal coordination is disrupted. SC formation may initiate on incompletely replicated chromosomes, creating the potential for replication forks to collide with SC components—causing replication stress and DNA damage. Alternatively, Atx2 may be required for a checkpoint that delays SC assembly until replication is complete. Clarifying why oocytes are particularly sensitive to such disruptions remains an important area for future investigation.

Although the mechanisms regulating premeiotic DNA replication in metazoans are not well defined, recent findings in mice highlight its importance. In female germ cells, delayed premeiotic replication leads to premature oocyte loss and increased DNA damage, as shown by elevated DNA damage response markers and TUNEL staining—paralleling our findings in Drosophila. These similarities suggest that while the specific molecular players may differ, precise timing of premeiotic DNA replication is an evolutionarily conserved requirement for maintaining oocyte genome integrity and fertility.

In summary, our work reveals a previously unrecognized role for the conserved RBP Atx2 in orchestrating the meiotic entry. By ensuring timely premeiotic DNA replication and maintaining genome stability, Atx2 is essential for meiotic progression and oocyte viability. These findings uncover a critical function for Atx2 in germline development and highlight the broader importance of post-transcriptional regulation in meiosis initiation.

## ACKNOWLEDGMENTS

We thank Rui Martinho, Andrew Swan, Craig Smibert, and Prashanth Rangan for helpful experimental suggestions and comments on the manuscript. We thank Bloomington Drosophila Stock Center (NIH P40 OD018537), Mary Lilly, Liz Gavis and R. Scott Hawley for Drosophila stocks. We observed the 1B1, orb and γH2Av monoclonal antibodies developed by H.D. Lipshitz, P. Schedl and R.S. Hawley, respectively, from the Developmental Studies Hybridoma Bank, created by the NICHD of the NIH and maintained at The University of Iowa, Department of Biology, Iowa City, IA 52242. This work was supported by the Canadian Institutes of Health Research (FRN 159510) and T.R.H. is part of the University of Toronto Medicine by Design initiative, which receives funding from CFREF. V.L.M. is supported by an Ontario Graduate Scholarship. C.R. is supported by an Irish Research Council Postgraduate Award and B.B. by a Wellcome-DBT IA Intermediate Fellowship, and M.R. by grants from Science Foundation Ireland and a Research Ireland Future Frontiers Awards.

## AUTHOR CONTRIBUTIONS

Conceptualization, V.L.M., and T.R.H.; methodology, V.L.M., and C.Y; reagents, V.L.M., C.R., B.B, and M.R.; investigation, V.L.M, and T.R.H.; writing – original draft, V.L.M, and T.R.H; writing – review & editing, V.L.M, T.R.H, C.Y., C.R, B.B., and M.R.; funding acquisition, V.L.M., and T.R.H.; supervision, T.R.H.

## MATERIALS AND METHODS

### Drosophila Husbandry

All fly stocks were reared at 25 °C with controlled humidity on standard medium (cornmeal, agar, yeast and molasses). All stocks used are listed in Table S1. Genotypes for each figure are listed in Table S2. All knockdown and mutant experiments were conducted at 25 °C except for *dap* and *dap^CRL4^* overexpression which was done at 29 °C.

### Generation of endogenous FRT-flanked Atx2::GFP

CRISPR genome engineering method^88^ with homologous recombination (HR) was used to generate FRT-flanked Atx2::GFP line. PUC19-Atx2::GFP vector, as described in Bakthavachalu et al. 2018, was used as a template, this plasmid already contained an FRT site at the 3’UTR of the *Atx2* locus. A fragment containing 5’ UTR FRT was synthetized by BioCat company, amplified by PCR (5’-ttgaatacaaaccataagcagttcaaattg-3’; 5’ aacaacaacaattgcattcattttatgttt-3’) and inserted in the open PUC19 Atx2 plasmid through Gibson cloning^89^. To facilitate screening, a dsRed protein driven by 3XP3 eye promoter was inserted in the first intron of PUC19 Atx2 plasmid, transcribed in opposite orientation to *Atx2* gene. The pU6 dual gRNA (5’-TTTGTGTCCGATTGCCGGG-3’ and 5’-GGAATGGCGCACGGGAAGG-3’) containing plasmid was mixed with PUC19 FRT-flanked Atx2::GFP plasmid in 1:3 ratios and injected into *act5C-Cas9* flies and screened for dsRed eye marker (BestGene Inc.). The established fly lines were then sequence validated.

### Generation of UASz-Atx2 isoforms

The *UASz-3xHA::Atx2-B* and *UASz-3xHA::Atx2-D* plasmids were generated using Q5 High-Fidelity DNA Polymerase (NEB, M0491) and 2x GB-AMP PaCeR HP Master Mix (GeneBio Systems) for PCR amplification and NEBuilder HiFi DNA Assembly Master Mix (NEB, E2621) for plasmid assembly into pUASz^90^. Atx2-B and Atx2-D isoforms were PCR amplified from pAc-3xFlag-Atx2 plasmid (gift from Craig Smibert) as two fragments to mutate the site targeted by HMS01392 shRNA carried by the BDSC stock #36114. The 21-bp target site was mutated from CC<u>G</u>GA<u>C</u>GA<u>G</u>AA<u>G</u>GA<u>ATTG</u>GA<u>A</u> to CC<u>c</u>GAtGA<u>a</u>AA<u>a</u>GA<u>gctc</u>GA<u>g</u> through PCR. A 3xHA fragment was added to the N-terminus of each construct. The UASz constructs were injected into the attP40 landing site in the *yw, P{nos-phiC31\int.NLS}X; P{CaryP}attP40* strain by Rainbow Transgenic Flies, Inc.

### Egg lay, dorsal appendages and hatching

Six 2-3 days old females of the respective genotypes and four *w^11^*^18^ males were maintained on grape juice plates with yeast paste. Fresh grape juice plates with yeast paste were exchanged every 24 hours for 3 days. The number of eggs laid were counted and averaged over the three days. For hatching, hatched eggs on grape juice plates were counted 24 - 48 hours after egg lays. To evaluate the dorsal appendages, the eggs laid by six females on grape juice plates with yeast paste were scored based on phenotype: wild type – two separate dorsal appendages, moderately ventralized – fused or partially fused dorsal appendages, and ventralized – no dorsal appendages visible. Representative images were taken with a Nikon DS-Fi1-L2 camera.

### Ovary immunofluorescence

Adult ovaries were stained according to standard procedures. Briefly, ovaries from yeasted flies were dissected in PBS and fixed in 4% formaldehyde (Thermo Scientific, 28908) in PBS for 15 min. Ovaries were then permeabilized with 1% Triton X-100 (BioShop Canada, TRX506) in PBS for 20 min. Ovaries were incubated with primary antibodies diluted in 1% PBST (1% (w/v) bovine serum albumin [BSA, BioShop Canada, ALB001], 0.1% Triton X-100, PBS) overnight at 4°C followed by incubation with the appropriate secondary antibodies diluted in 1% PBST for 2 hours at room temperature. Ovaries were counterstained with 4′,6-diamidino-2-phenylindole (0.2 – 1 µg/mL; Cell Signaling Technologies, 4083S) for 15 min and/or phalloidin Alexa Fluor 633 (1:250; Invitrogen, A22284) for 2 hours with secondary antibodies. Ovaries were mounted in VECTASHIELD Antifade Mounting Medium (BioLynx, VECTH1000). All images were acquired with a Leica SP8 inverted scanning confocal microscope using 63x (NA 1.4, immersion oil) objectives. All experiments were performed using multiple sections (z-stacks) from confocal images. Image analysis and maximum projections were performed using FIJI^91^. All confocal images shown are maximum projections of 3 slices at 0.3 µm each.

The following primary antibodies were used: mouse anti-1B1 (1:50; deposited to the DSHB by H.D. Lipshitz); mouse IgG1 anti-orb 4H8 (1:100; deposited to the DSHB by P. Schedl); mouse IgG2b anti-γH2Av (1:5000; UNC93-5.2.1 deposited to DSHB by R.S. Hawley); mouse IgG1 anti-C(3)G (1:500; a gift from R.S. Hawley); and rabbit anti-GFP (1:1000; Invitrogen, A11122).

The following secondary antibodies were used at a 1:500 dilution: donkey anti-mouse Cy3 (Jackson ImmunoResearch Labs, 715-165-151); donkey anti-rabbit Alexa Fluor 647 (Jackson ImmunoResearch Labs, 711-605-152); goat anti-rabbit Oregon Green 488 (Invitrogen, O-11038); goat anti-mouse Alexa Fluor 488 (Invitrogen, A-11001); goat anti-mouse IgG1 Alexa Fluor 488 (Invitrogen, A-21121); and goat anti-mouse IgG2b Alexa Fluor 568 (Invitrogen, A-21144).

### EdU incorporation assay

EdU incorporation assay was performed using Click-iT EdU Cell Proliferation Kit for Imaging, Alexa Fluor 647 dye (Invitrogen, C10340) and according to manufacturer’s protocol. Briefly, adult ovaries from yeasted flies were dissected in PBS and incubated in 10 µm of EdU in Schneider’s Drosophila Medium (Gibco, 21720024) with 10% fetal bovine serum (FBS; Gibco, 12483020) for 1 hour at room temperature. Medium was removed and ovaries were washed twice with 10% FBS in Schneider’s Drosophila Medium for 3 min each. Ovaries were then fixed in 4% formaldehyde in PBS for 20 min. Ovaries were subsequently washed twice with 3% BSA in PBS for 5 min before permeabilization with 0.5% Triton X-100 in PBS for 20 min followed by two washes with 3% BSA in PBS. Ovaries were incubated in the dark at room temperature with the Click-iT reaction cocktail exactly according to manufacturer’s protocol. Following Click-iT reaction, ovaries were washed twice 3% BSA in PBS, and primary and secondary antibody staining was done according to standard ovary immunofluorescence protocol in the dark.

### Quantification of karyosome morphology

Oocyte karyosomes were analyzed in stage 4-5 egg chambers. They were scored as spherical if all DNA was condensed and round, mildly distorted if DNA was almost completely condensed but not visibly round, and distorted if DNA was not condensed and spread apart.

### RNA extraction and RT-qPCR

Total RNA was extracted from three yeasted ovaries according to manufacturer’s protocol using TRI-Reagent, chloroform (Sigma-Aldrich, 472476), GlycoBlue (ThermoFisher Scientific, AM9516), 2-propanol (Sigma-Aldrich, I9516) and ethanol, and eluted in nuclease-free water. Purified RNA was treated with Turbo DNAse (ThermoFisher Scientific, 2238G2). For cDNA synthesis, 2000 ng of RNA was used with SuperScript IV Reverse Transcriptase (ThermoFisher Scientific, 18090050) and oligo dT-20mer (Integrated DNA Technology, 51-01-15-01).

Quantitative PCRs was carried out on 1/50 of reverse transcription reactions using the SensiFAST SYBR No-ROX kit (FroggaBio, BIO-98050) and a Bio-Rad CFX384/C1000 Touch system (Bio-Rad). The PCR program was as follows: 2 min at 95 °C; 45 cycles of 95 °C for 5 s and 60 °C for 30 s. Results of qPCRs for transposable elements were normalized to the mean of value obtained of CG8187, CG2698 and Und. Results were calculated using the ΔΔCt method: ΔΔCt = 2^-(ΔCt*^Atx2^* ^RNAi^ – ΔCt*^mCherry^* ^RNAi^), where ΔCt = Ct (gene) – Ct (mean of CG8187, CG2698 and Und). Primers used are listed in Table S3.

### Statistics and reproducibility

Graphs and statistics in relevant figures were generated using Prism 10.1 (GraphPad) or in R. All statistical analysis performed is noted in the figure captions. All fly crosses were repeated at least once and over 10 flies were examined for each experiment. Matched replicates in each graph are colour coded, and the number of germaria/oocytes examined for each replicate is indicated in the figure caption.

---

**Figure S1.**
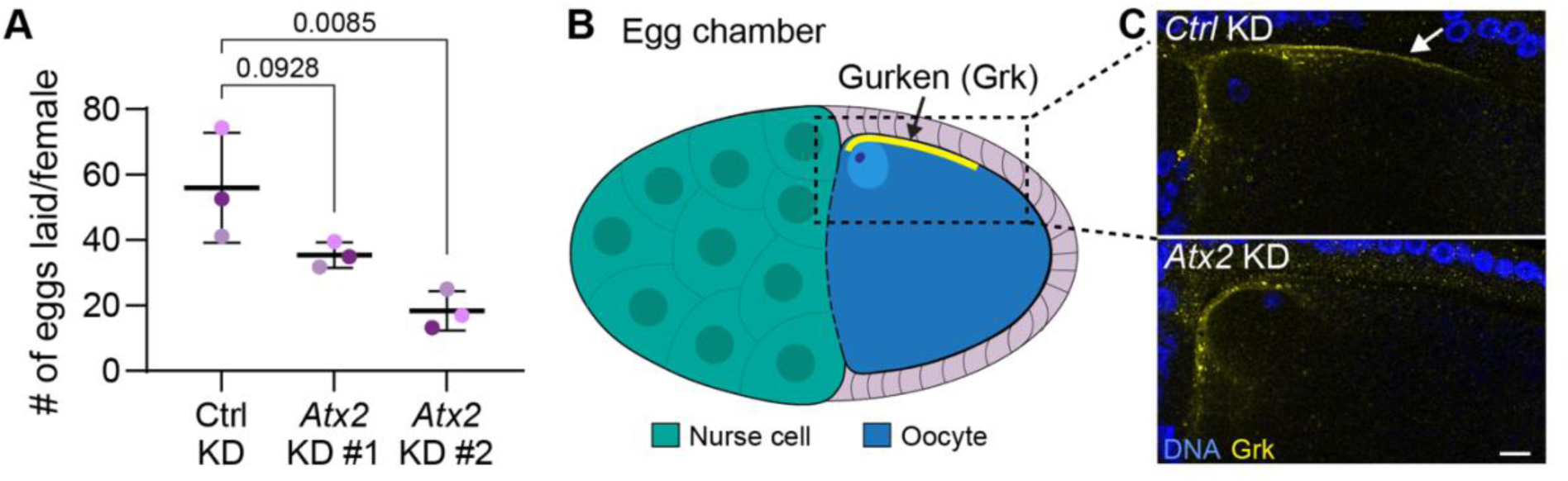
Atx2 depletion reduces egg laying and oocyte Gurken protein levels. **(A)** Number of eggs laid per female expressing Ctrl (*mCherry*) shRNA or *Atx2* shRNA. Data are means ± s.d. of three independent replicates (>100 eggs per replicate). *p*-values, one-way ANOVA followed by Dunnett’s multiple comparison test. **(B)** Left, graphical representation of a stage 10 egg chamber expressing Grk (yellow) along the dorsal-anterior side of the oocyte. Right, representative confocal micrographs of stage 10 egg chambers of females expressing Ctrl (*mCherry*) or *Atx2* shRNA in the germline stained with anti-Grk (yellow) and DAPI (blue). Scale bar, 10 µm.

**Figure S2.**
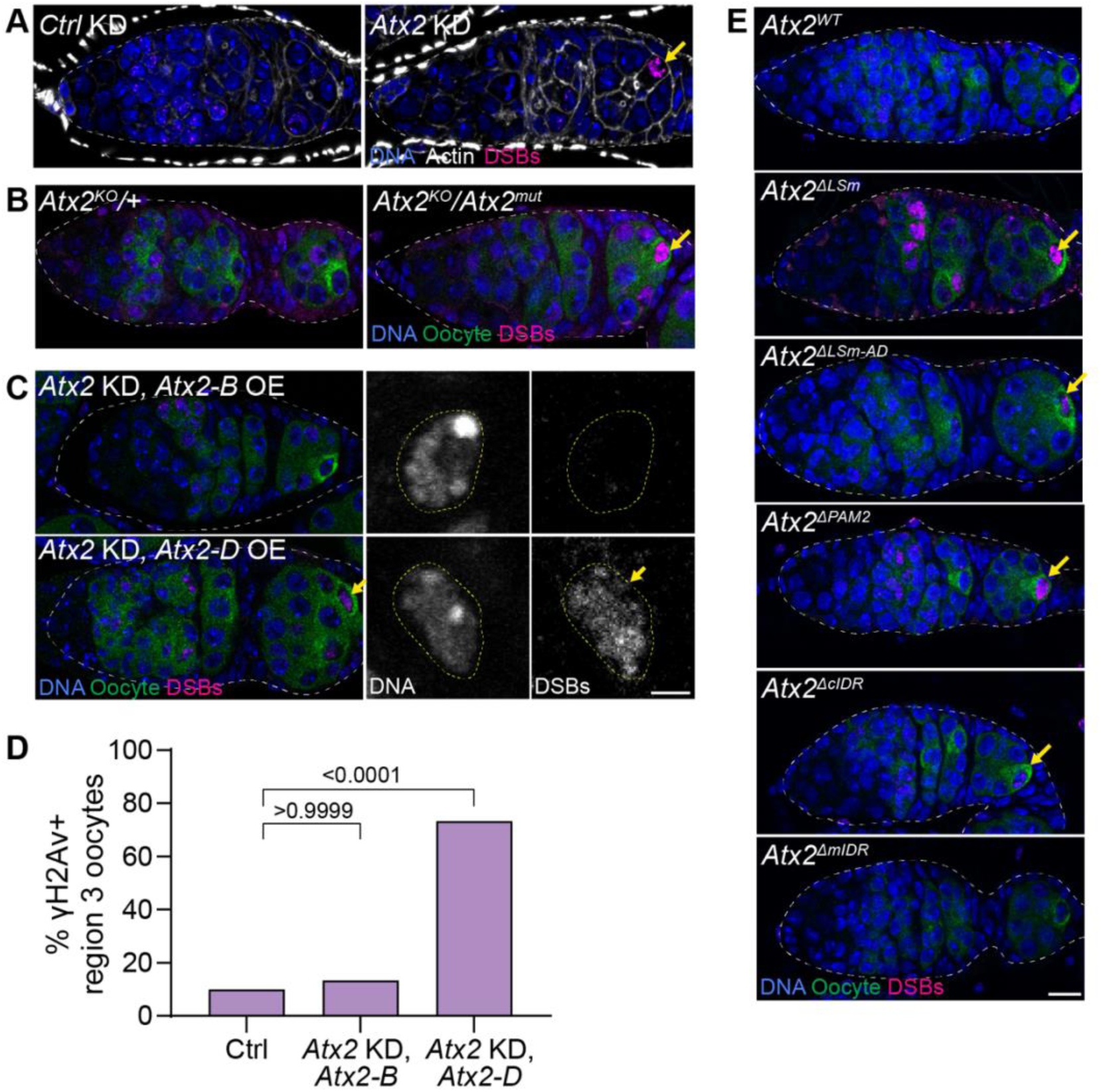
*Atx2-B* post-transcriptionally maintains oocyte genome stability during meiosis. **(A)** Confocal micrographs of germaria expressing Ctrl (*mCherry*) or *Atx2* shRNA in the germline stained with anti-γH2Av (DSBs, magenta), anti-Orb (oocyte, green) and DAPI (DNA, blue) Yellow arrows indicated persistent γH2Av staining. Scale bars, 10 µm. **(B)** Confocal micrographs of germaria from heterozygous control and germline ablated *Atx2* ovaries stained with anti-γH2Av (DSBs, magenta), anti-Orb (oocyte, green) and DAPI (DNA, blue). Yellow arrows indicated persistent γH2Av staining. Scale bars, 10 µm. **(C)** Confocal micrographs of germaria (left) and region 3 oocytes (right) expressing shRNA resistant *Atx2-B* or *Atx2-D* in the *Atx2* shRNA knockdown germlines stained with anti-γH2Av (DSBs, magenta), anti-Orb (oocyte, green) and DAPI (DNA, blue). Yellow arrows indicated persistent γH2Av staining. Scale bars, 10 µm (germaria) and 2 µm (oocytes). **(D)** Quantification of (**C**). Percent of γH2Av positive region 3 oocytes of indicated genotypes (n = 30 for each genotype). *p*-values, Fisher’s exact test. **(E)** Confocal micrographs of germaria expressing full-length *Atx2*, *Atx2^ΔLSm^*, *Atx2^ΔLSm-AD^*, *Atx2^ΔPAM2^*, *Atx2^ΔmIDR^* and *Atx2^ΔcIDR^* under native regulatory elements in *Atx2* germline ablated ovaries anti-γH2Av (DSBs, magenta), anti-Orb (oocyte, green) and DAPI (DNA, blue). Yellow arrows indicated persistent γH2Av staining. Scale bars, 10 µm.

**Figure S3.**
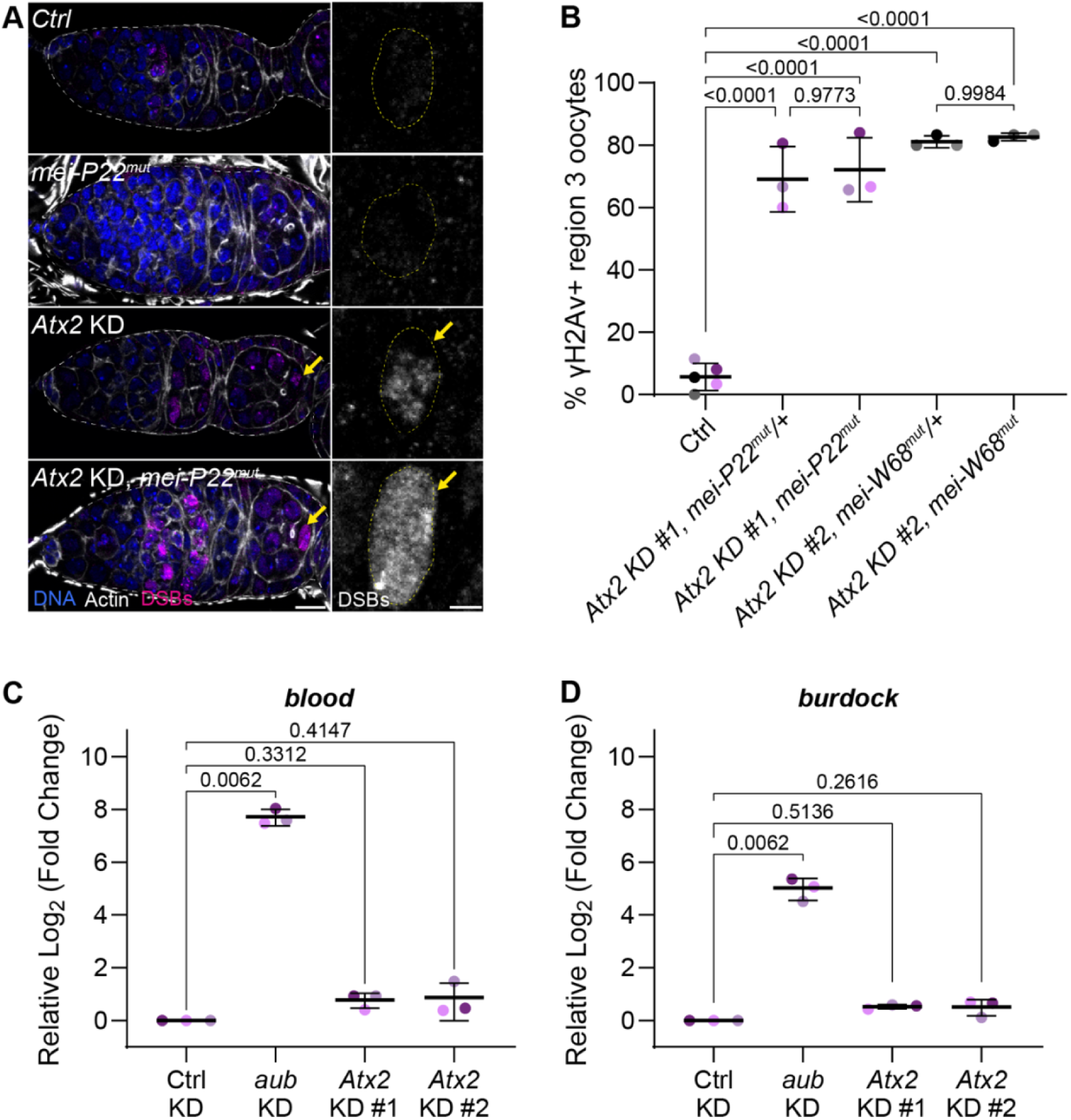
Oocyte DNA damage occurs independently of meiotic DSBs and transposons in *Atx2* knockdowns. **(A)** Confocal micrographs of germaria (left) and region 3 oocytes (right) from Ctrl, *mei-P22^mut^*, germline *Atx2* shRNA, and *mei-P22^mut^* and germline *Atx2* shRNA stained with anti-γH2Av (DSBs, magenta), phalloidin (actin, white) and DAPI (DNA, blue). Yellow arrows indicated persistent γH2Av staining. Scale bars, 10 µm (germaria) and 2 µm (oocytes). **(B)** Quantification of **(A)**. Percent γH2Av positive region 3 oocytes of the indicated genotypes. Data are mean ± s.d. (10 – 37 oocytes per replicate). *p*-value, ANOVA, Tukey’s multiple comparison test. **(C, D)** Relative fold change of *blood* **(C)** and *burdock* **(D)** long terminal repeat elements in ovaries expressing Ctrl (*mCherry*), *aub*, *Atx2* shRNAs in the germline. Data are means ± s.d. *p*-values, Kruskal-Wallis, Dunn’s multiple comparison test.

**Figure S4.**
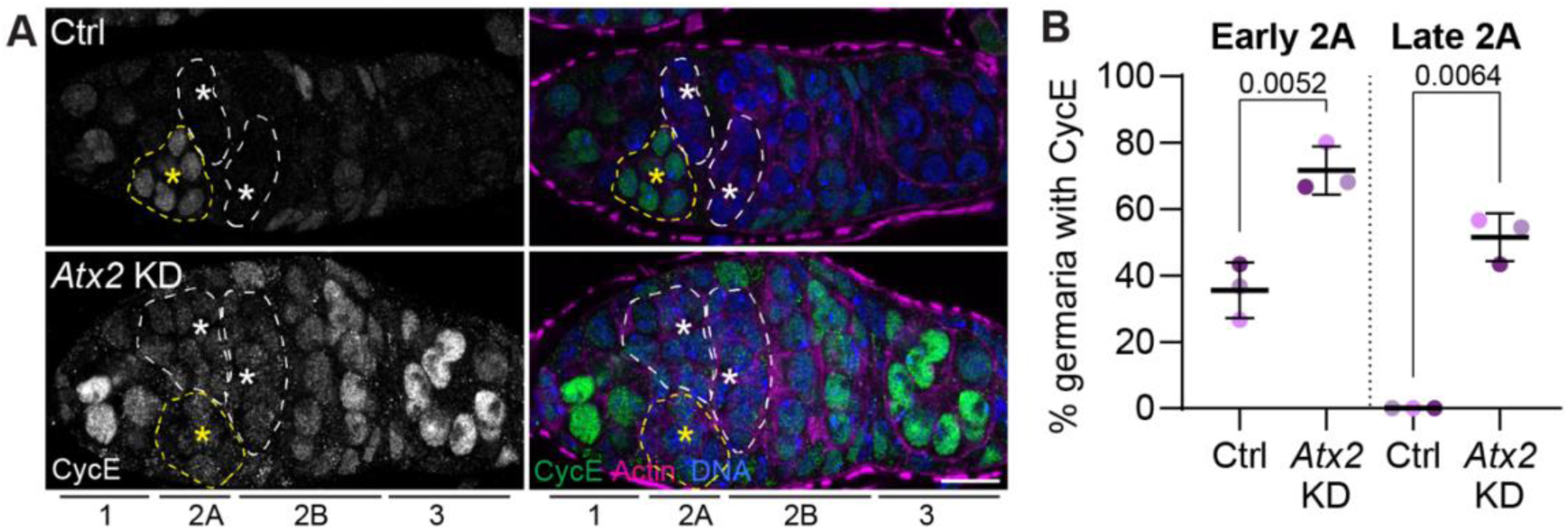
CycE levels remain elevated in *Atx2* knockdowns. **(A)** Confocal micrographs of CycE-GFP germaria expressing Ctrl (*LexA*) or *Atx2* shRNAs in the germline stained with anti-GFP (Dap, green), anti-1B1 (soma/fusome, magenta) and DAPI (DNA, blue). Scale bar, 10 µm. Yellow dashed line, early region 2A. While dashed line, region 2B. **(B)** Quantification of **(A)**. Percent of germaria with above background levels of CycE-GFP in the early or late region 2A cysts of the indicated genotypes. Data are means ± s.d. (22 – 30 germaria per replicate). *p*-value, two-tailed Welch’s t test.

**Figure S5.**
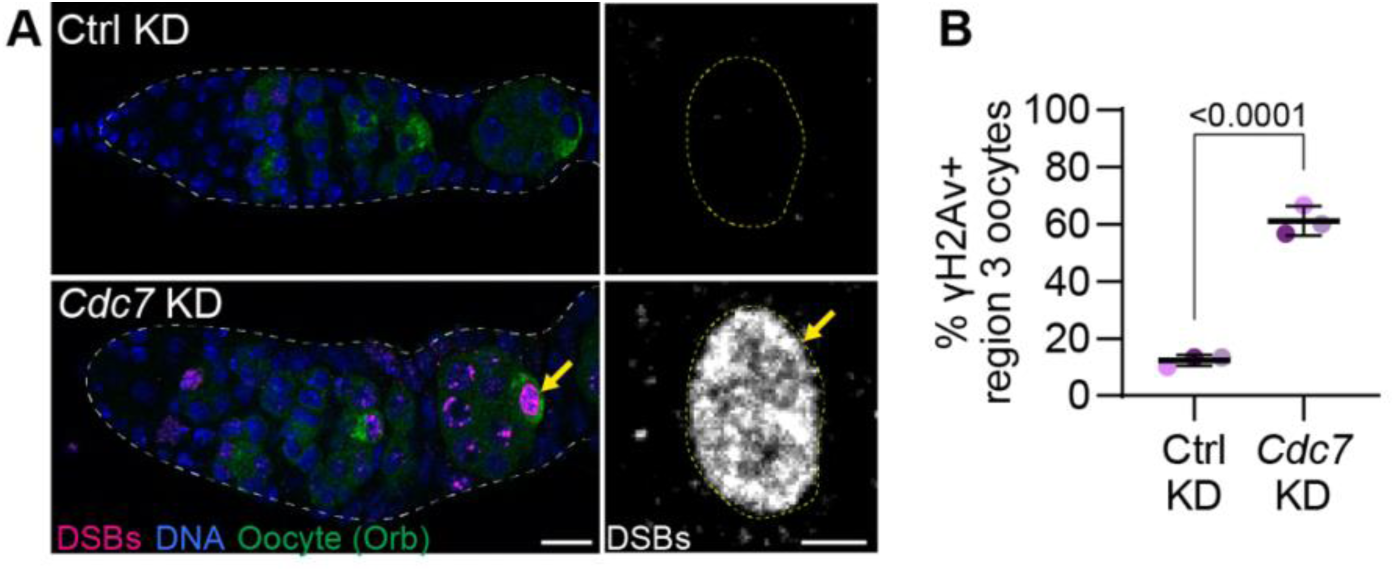
*Cdc7* is required to maintain oocyte genome stability. **(A)** Confocal micrographs of germaria (left) and region 3 oocytes (right) expressing Ctrl (*mCherry*) or *Cdc7* shRNA in the germline stained with anti-γH2Av (DSB, magenta), anti-Orb (oocyte, green) and DAPI (DNA, blue). Scale bar, 10 µm (germaria) and 2 µm (oocyte). White dashed line, germline. Yellow dashed line, oocyte nucleus. **(B)** Quantification of **(A)**. Percent of γH2Av positive region 3 oocytes of indicated genotypes. Data represent means ± s.d. of three independent replicates (12 – 30 oocytes per-replicate). *p*-values, unpaired t-test.

**Figure S6.**
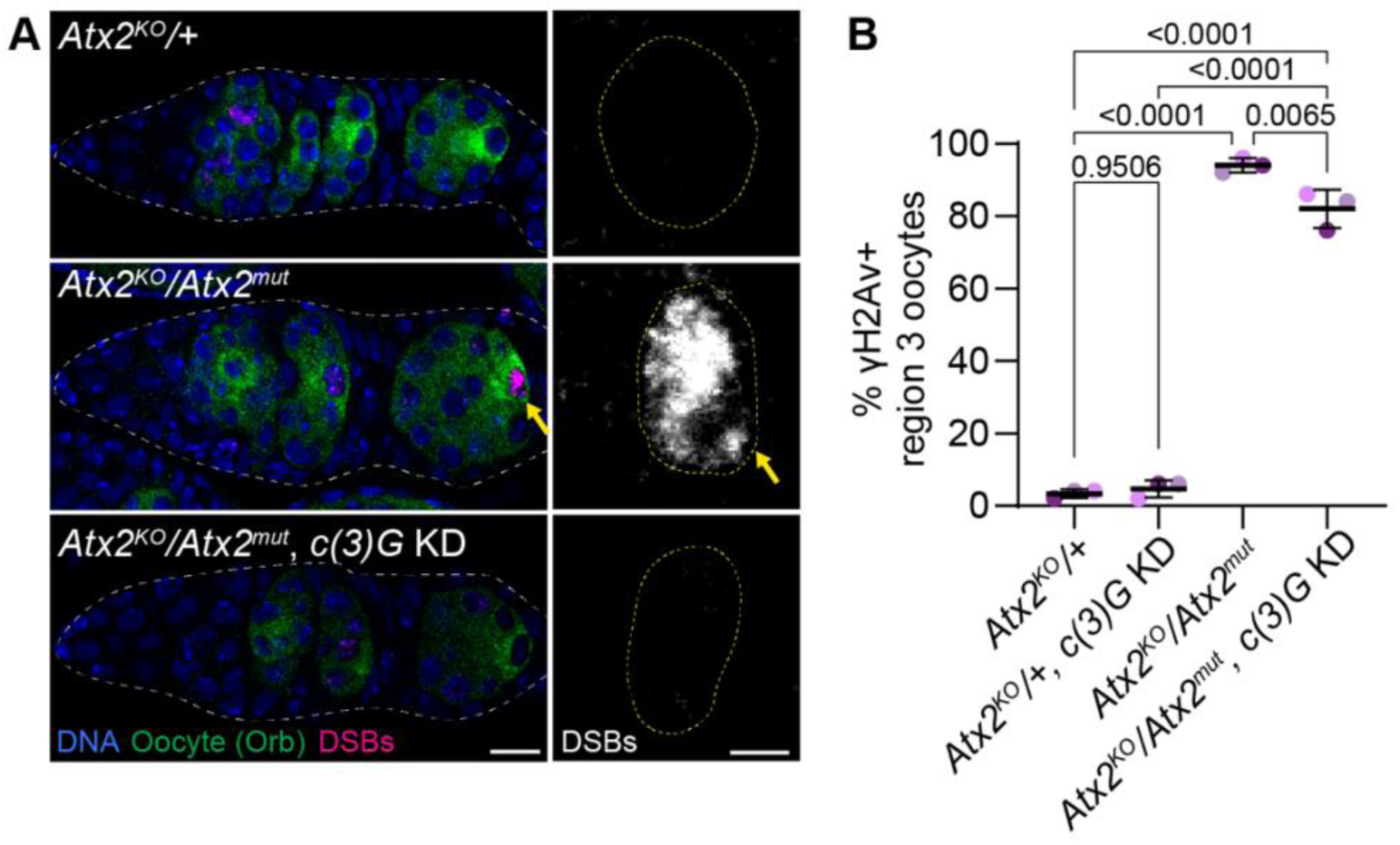
Synaptonemal complex knockdown reduces frequency of oocyte genome instability in *Atx2* knockouts. (A) Confocal micrographs of germaria (left) and region 3 oocytes (right) of the following genotypes: *Atx2^KO^/+* heterozygotes; *Atx2* germline ablated *Atx2^KO^/Atx2^mut^* transheterozygotes; and *c(3)G* shRNA expressed in the germlines of *Atx2* germline ablated *Atx2^KO^/Atx2^mut^*transheterozygotes. Ovaries were stained with anti-γH2Av (DSB, magenta), anti-Orb (oocyte, green) and DAPI (DNA, blue). Scale bar, 10 µm (germaria) and 2 µm (oocyte). White dashed line, germline. Yellow dashed line, oocyte nucleus. Yellow arrow, persistent oocyte γH2Av. (B) Quantification of **(A)**. Percentage of γH2Av positive region 3 oocytes of the indicated genotypes. Data are means ± s.d. of three independent replicates (50 oocytes per replicate). *p*-value, ANOVA, Tukey’s multiple comparison test.

**Table S1.**
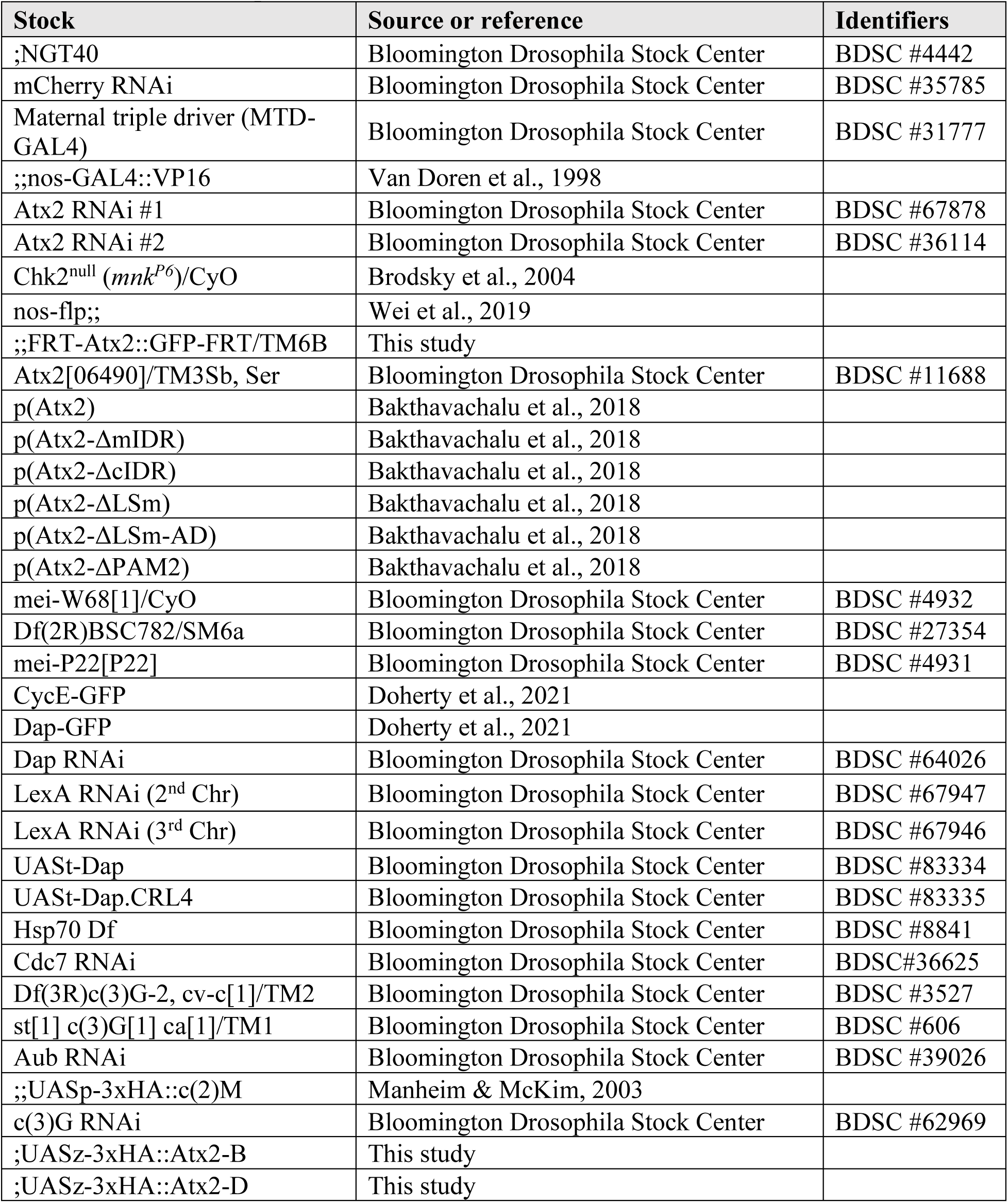
List of Drosophila stocks.

**Table S2.**
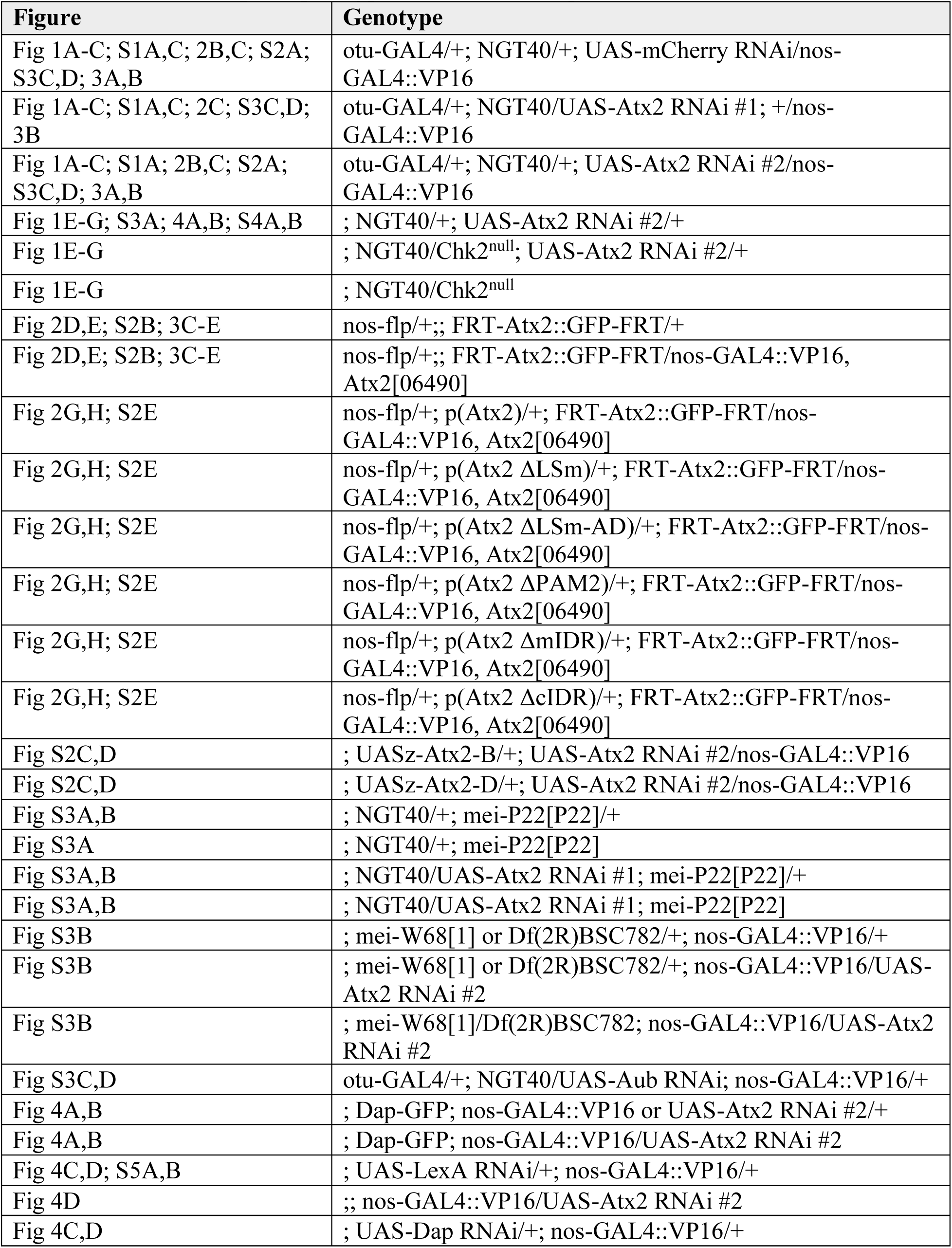
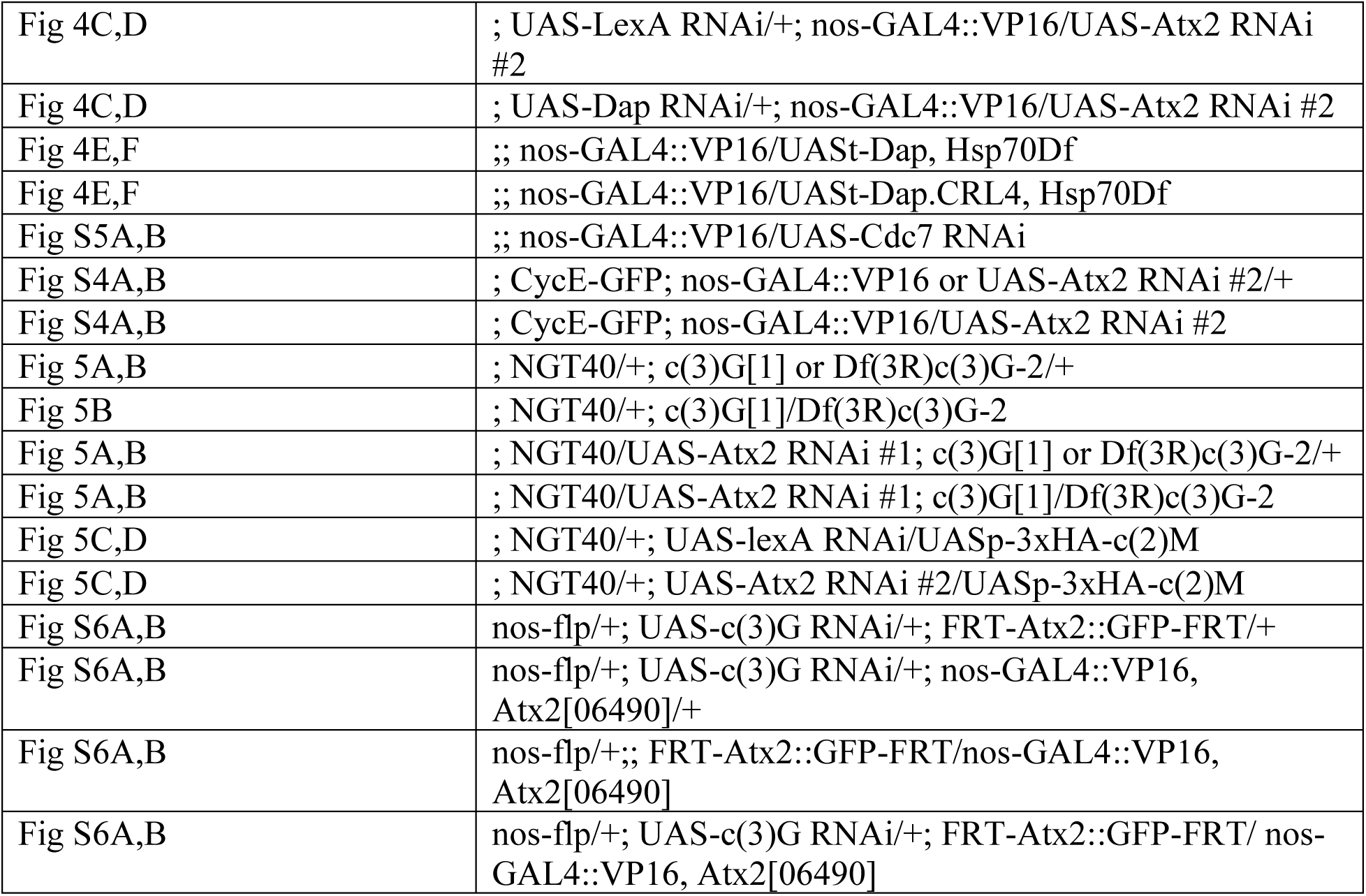
List of Drosophila genotypes used for each figure.

**Table S3.**
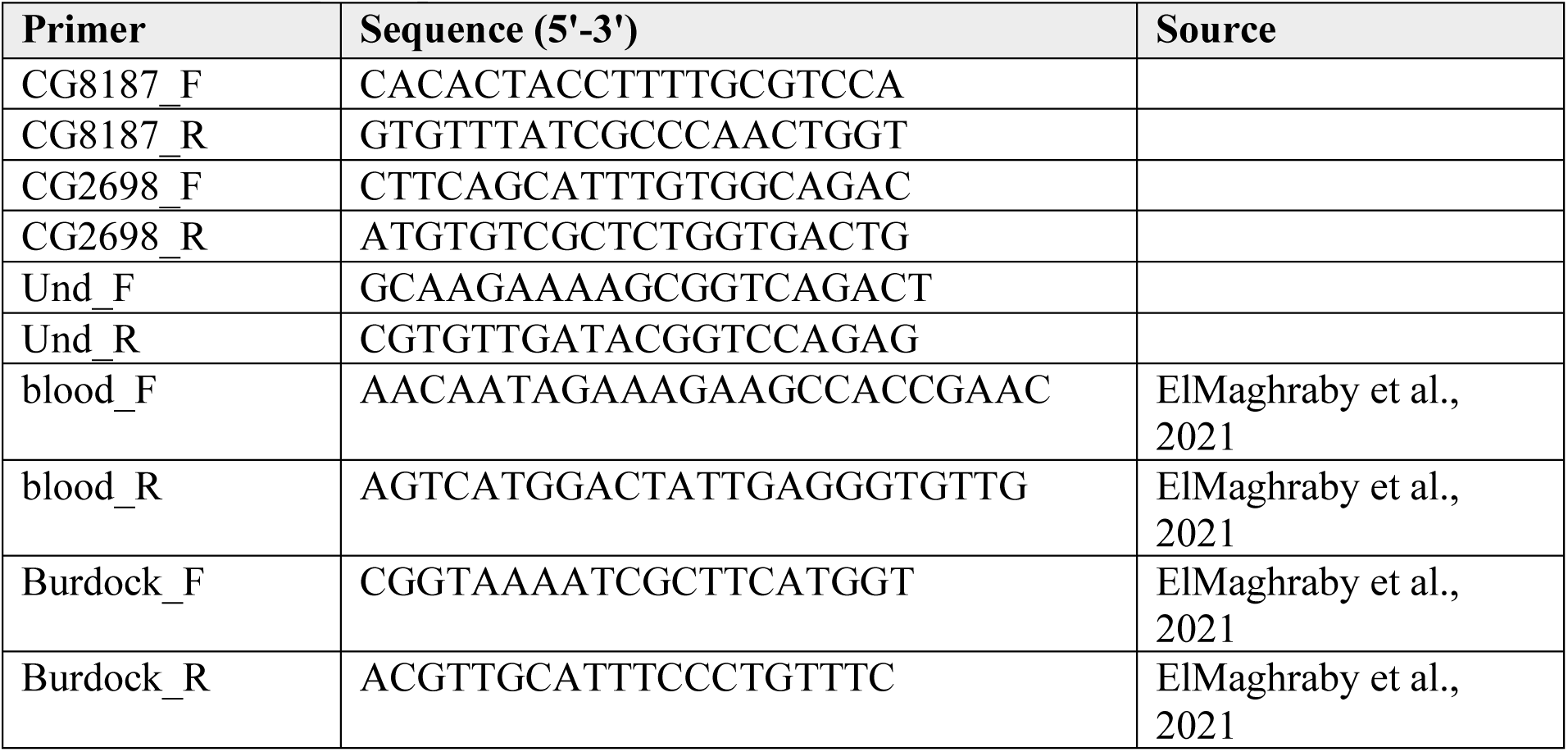
List of qPCR primers.

